# Dilp8 controls a time window for tissue size adjustment in *Drosophila*

**DOI:** 10.1101/2020.11.09.375063

**Authors:** L. Boulan, D. Blanco-Obregon, K. El Marzkioui, F. Brutscher, D.S. Andersen, J. Colombani, S. Narasimha, P. Léopold

## Abstract

The control of organ size mainly relies on precise autonomous growth programs. However, organ development is subject to random variations, called developmental noise, best revealed by the fluctuating asymmetry observed between bilateral organs. The developmental mechanisms ensuring bilateral symmetry in organ size are mostly unknown. In *Drosophila*, null mutations for the relaxin-like hormone Dilp8 increase wing fluctuating asymmetry, suggesting that Dilp8 plays a role in buffering developmental noise. Here we show that size adjustment of the wing primordia involves a peak of Dilp8 expression that takes place sharply at the end of juvenile growth. Wing size adjustment relies on a cross-organ communication involving the epidermis as the source of Dilp8. We identify ecdysone signaling as both the trigger for epidermal *dilp8* expression and its downstream target in the wing primordia, thereby establishing reciprocal feedback between the two hormones as a systemic mechanism controlling organ size precision. Our results reveal a hormone-based time window ensuring fine-tuning of organ size and bilateral symmetry.

## INTRODUCTION

One striking aspect of developmental processes is the precision with which final organ size is achieved and coordinated with other organs’ dimensions to give rise to individuals with adequate proportions and functions. Although many developmental processes are now being characterized with great detail, the mechanisms of determination and fine adjustment of organ size are not understood correctly.

Symmetric bilateral organs constitute an ideal model for the study of developmental precision^1,2^. Since in most cases, left and right bilateral organs develop in the same environment, the limited, random asymmetries observed in adult bilateral traits illustrates the stochastic variations taking place during development, also called developmental noise^3^. Developmental noise is generally evaluated through a measure of the fluctuating asymmetry (FA) index, calculated as the variance of the mean-scaled difference between left and right bilateral traits^1,2^. The low levels of variability observed between bilateral organs in physiological conditions suggest that buffering mechanisms are at play that maintain developmental robustness^4^, although this is still under debate^5,6^.

The identification of mutations affecting developmental precision in genetically tractable models opens the possibility to address such buffering mechanisms^7^–^10^. In *Drosophila*, mutations in the relaxin-like hormone Dilp8 and its receptor Lgr3 were recently found to decrease developmental stability through a systemic relay. Dilp8 was first identified as a signal produced by injured or tumorous imaginal discs that induces a delay in development allowing for tissue repair^11,12^. This delay is mediated by the regulation of the steroid hormone ecdysone through a neural circuitry involving the Dilp8 receptor Lgr3^13^–^15^. In the absence of tissue injury, the removal of Dilp8 or Lgr3 function increases FA in adult wings, indicating a physiological role for the Dip8/Lgr3 axis in the control of developmental stability^12^–^15^.

Although the mechanism of tissue repair-induced delay by Dilp8 is now better understood^16^, the mechanism by which the Dilp8 hormone controls developmental stability remains unknown. Two distinct hypotheses could account for such control. Continuous feedbacks taking place during the growth phase could maintain organs on a growth trajectory leading to an appropriate final size. In that case, a robustness factor would be expected to come from the organ itself as part of the feedback mechanism. Alternatively, developing organs could randomly deviate from a standard growth trajectory up to a time window in development where the extent of the deviation is evaluated and a correction is made. If so, robustness factors could control either the emergence of the time window, the measure of the deviation, or its correction.

To address these hypotheses experimentally, we first quantified the size variations of wing discs pairs along larval and early pupal development. We found that FA, while elevated during the growing larval stage, is sharply corrected after the larval-to-pupa (L/P) transition. In line with this finding, we observed that Dilp8 expression is strongly upregulated in the epidermis and functionally required at the L/P transition for maintaining low FA. We also established that the sharp burst of epidermal Dilp8 expression is directly controlled by the rise of ecdysone titer at the end of larval development. Finally, our results indicate that Dilp8 is in turn required to adjust the levels of ecdysone at the L/P transition.

We therefore propose a model whereby developmental stability is ensured through a hormonal cross-talk between ecdysone and Dilp8, establishing a precise developmental time window past which fluctuating asymmetry, an indicator of size adjustment, is buffered.

## RESULTS

### A time window for wing imaginal disc size adjustment during development

As a first approach, and to distinguish between a “feedback” and a “time window” mode of size adjustment, we aimed to establish when the size of paired organs is adjusted by quantifying the left-right differences and FA in wing imaginal discs during development. Volume quantification was performed after 3D reconstruction of the GFP-labelled *nubbin* expression domain of wing imaginal discs (*nub>GFP*), the so-called wing “pouch” corresponding to the presumptive wing blade (Fig. 1a, see Methods). The left-right (L-R) volume difference was plotted for dissected pairs of discs at two timepoints during the growth phase (96h after egg deposition (AED), mid 3^rd^ larval instar; and 114h AED, at the end of the 3^rd^ larval instar) and shortly after the larva-to-pupa (L/P) transition (7h after puparium formation (APF), after disc eversion and when dorsal and ventral sides of the presumptive wings have fully apposed). In control *dilp8^KO/+^* heterozygous animals, which display adult FA comparable to wild type animals^17^, the L-R variability of pouch volume is high at 96h AED with an FA index (FAi) around 40 (Fig. 1b,c; black dots and bars). The volume distribution is partially reduced at 114h AED, indicative of a first step in size adjustment taking place during larval development (FAi around 20). A major adjustment then occurs between 114h AED and 7h APF, with an FAi dropping to 10% of its value at 96h. The observation of a high FA during the growth phase, followed by a major correction around the L/P transition, suggests that a continuous feedback mechanism does not take place during development and favors a time window model.

**Figure 1.**
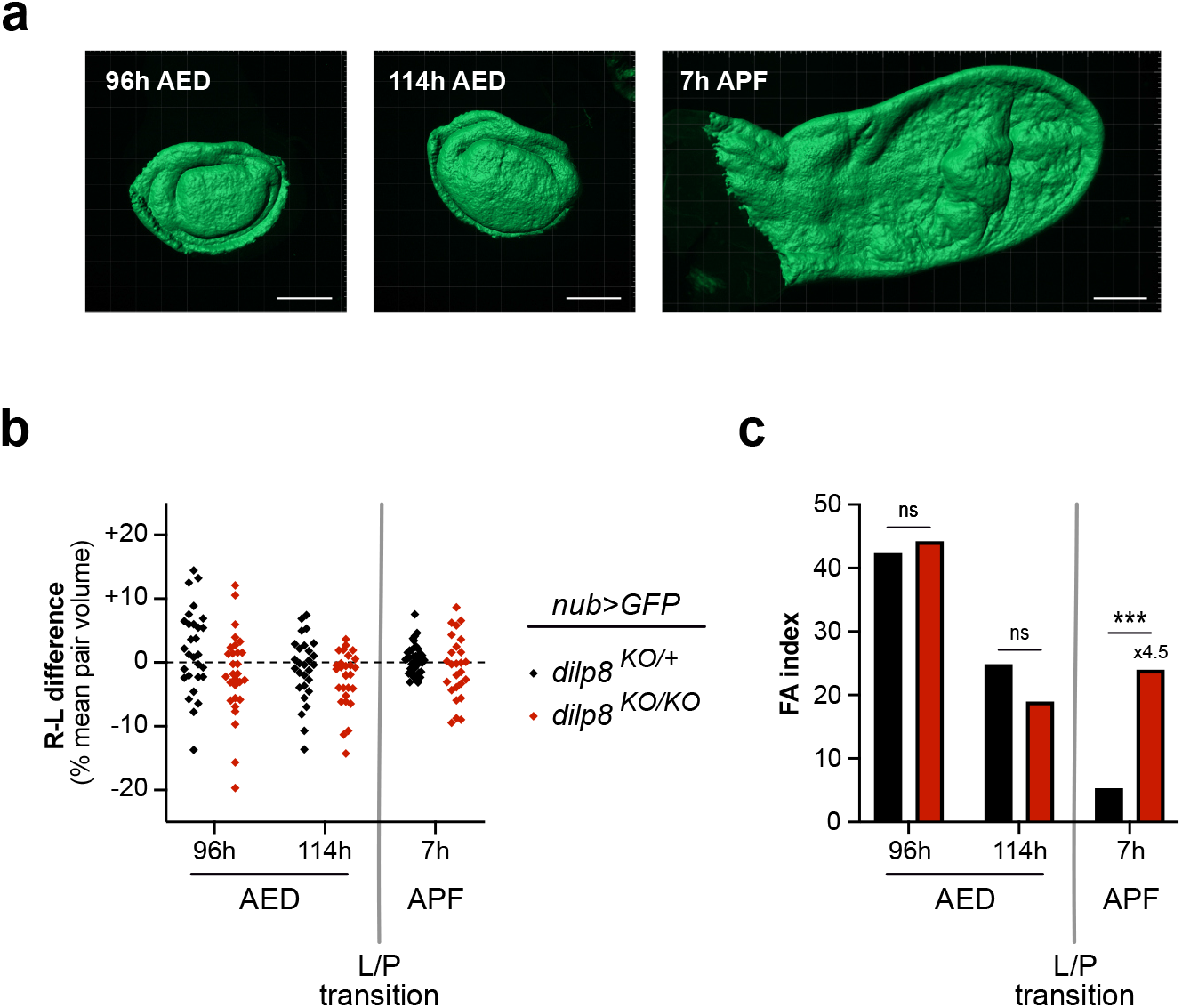
A developmental time window for organ size adjustment in early pupal development. (a) Representative examples of surface reconstruction for volume measurements of the wing pouch domain labelled with *nub>GFP* at 96h AED (mid L3 stage), 114h AED (late L3 stage) and 7h APF (early pupal stage). Scale bars represent 100 μm. (b) Distribution of the right-left (R-L) volume differences measured for individual pairs of wing discs and expressed as the percentage of the mean pair volume in control *dilp8* heterozygous animals (*dilp8^KO/+^, nub>GFP*) and null *dilp8* mutant animals (*dilp8^KO/KO^, nub>GFP*). (c) FA indexes calculated for each genotype and timepoint of the results shown in (b). At 96h AED, n=29 pairs of discs analyzed for each genotype; at 114h AED, n=29 for each genotype; at 7h APF, n=37 for *dilp8^KO/+^, nub>GFP* animals and n=26 for *dilp8^KO/KO^, nub>GFP* animals. *** p<0.001 and ns=not significant, F-tests. AED: after egg deposition, APF: after pupa formation, L/P transition: larva-to-pupa transition.

In order to understand the role of Dilp8 in buffering FA and determine when size adjustment is lost in the absence of Dilp8 function, we performed the same analysis for *dilp8^KO/KO^* null mutants. In this genetic context, while the initial drop in FAi was still present at 114h AED, there was a failure to resolve the L-R difference at the 7h APF time point (Fig. 1b,c; red dots and bars). Therefore, most of the difference in adult wing FA observed between *dilp8^KO/+^* and *dilp8^KO/KO^* (Fig. S1) is already established in early pupal development (7h APF, Fig. 1c). We conclude that Dilp8 is required for a major correction on wing disc size variability during a critical time window at the onset of pupal development.

### A pulse of Dilp8 expression at the larva-to-pupa transition controls organ size adjustment

Given the role of Dilp8 in setting a time window for size adjustment at the L/P transition, we precisely analyzed the timing of its expression around this transition. Using qRT-PCR on carefully staged animals, we observed that *dilp8* expression is kept at low basal levels during larval development and is sharply upregulated at a stage called white prepupa (WPP) marking the end of larval stage (Fig. 2a). This dramatic increase (1500-fold) in *dilp8* mRNA accumulation drops within 2h after WPP. Using the temperature-sensitive *GAL80^ts^* inhibitor combined with a ubiquitous *tub-GAL4 (tub^TS^>)* and a *dilp8-RNAi*, we downregulated *dilp8* at, and around, the WPP stage and analyzed adult wing FA. Abrogation of the *dilp8* expression peak around WPP induced an increase in wings FA comparable to constitutive *dilp8* inhibition (Fig. 2b). As a technical control, inducing a temperature shift silencing *dilp8* earlier during the larval L3 stage had no effect. Altogether, these results indicate that a peak of *dilp8* expression at WPP controls a time window for wing disc size adjustment occurring during early pupal development.

**Figure 2.**
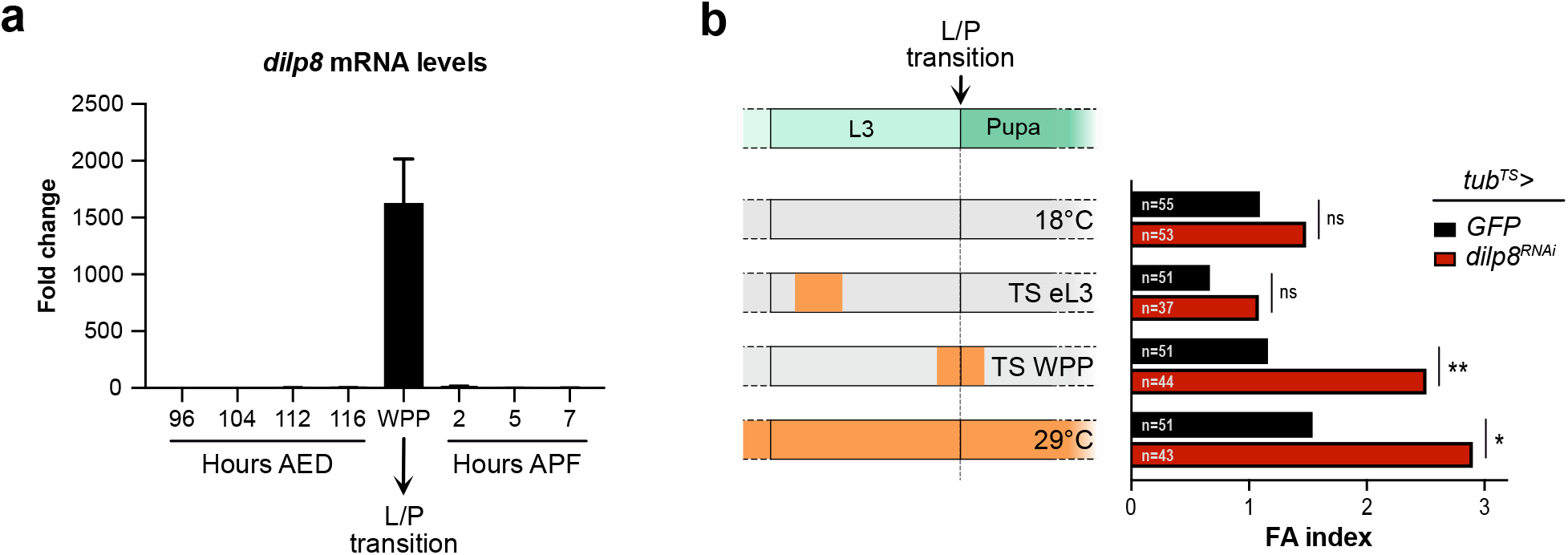
A peak of Dilp8 expression at the L/P transition is required for organ size adjustment. (a) Measurement of *dilp8* mRNA levels by qRT-PCR on whole animals (*w^1118^* control strain) at the indicated timepoints during development. Values are expressed as fold changes relative to the 96h AED timepoint. Error bars represent SEM. The white prepupa stage (WPP) corresponds to 0h APF and marks the L/P transition. (b) Temporal downregulation of *dilp8* using the ubiquitous *tub-GAL4, tub-GAL80^ts^ (tub^TS^>*) driver line crossed to a *UAS-dilp8^RNAi^* (TRIP) line or a control *UAS-GFP* line. The scheme on the left depicts the temperature-shift (TS) protocol. Orange periods mark the developmental times at which animals were switched from 18 to 29°C to activate the GAL4/UAS system and downregulate *dilp8.* Chronic 18°C and 29°C are the negative and positive controls, respectively. The graph on the right shows the FA indexes measured for adult pairs of wings of the given genotypes after the corresponding TS protocol. n values indicate the number of pairs analyzed; ** p<0.01, * p<0.05 and ns=not significant, F-tests. AED: after egg deposition, APF: after pupa formation, L/P transition: larva-to-pupa transition, WPP: white prepupa, eL3: early L3 stage.

### The larval epidermis is the source of Dilp8 for disc size adjustment

We next investigated the source of Dilp8 responsible for size adjustment during early pupal development. In the context of perturbed disc growth, Dilp8 is autonomously produced by the ill-growing discs and secreted into the hemolymph. However, inhibiting *dilp8* expression specifically in wing discs did not increase adult wing FA (Fig. S2a). This supports the notion that during normal growth, Dilp8-mediated size adjustment is not operating through a feedback mechanism where Dilp8 would be produced by the organ itself. To identify the source of *dilp8*, qRT-PCR was performed on dissected tissues at the WPP stage. While very low or no expression was detected in wing discs, fat body, gut, brain and salivary glands, high *dilp8* expression was detected in the carcass, mainly composed of epidermis and muscles apposed together (Fig. 3a). Co-immunostaining with muscle and epidermal markers in the context of a *dilp8-GFP* transcriptional reporter (see Methods and ^12^) indicated that *dilp8* is expressed specifically in the epidermis at the WPP stage (Fig. 3b,b). This result was confirmed using a *dilp8-lacZ^17^* reporter construct (Fig. S2b). In addition, the *dilp8-GFP* reporter showed a temporal upregulation at the WPP stage (Fig. S2c), in accordance with our expression data on whole animals. To confirm the epidermal origin of Dilp8 at the WPP stage, we silenced *dilp8* expression using two epidermal drivers (*Eip71CD-GAL4* and *E22C-GAL4*) and two separate *UAS-dilp8^RNAi^* lines. In these conditions, the quantification of *dilp8* mRNA levels on whole animals showed an abrogation of the peak of *dilp8* expression at the WPP stage (Fig. 3c). By contrast, silencing *dilp8* with two muscle-specific drivers failed to suppress *dilp8* expression at WPP (Fig. S2d), indicating that the larval epidermis is the unique source of *dilp8* expression at the WPP stage.

**Figure 3.**
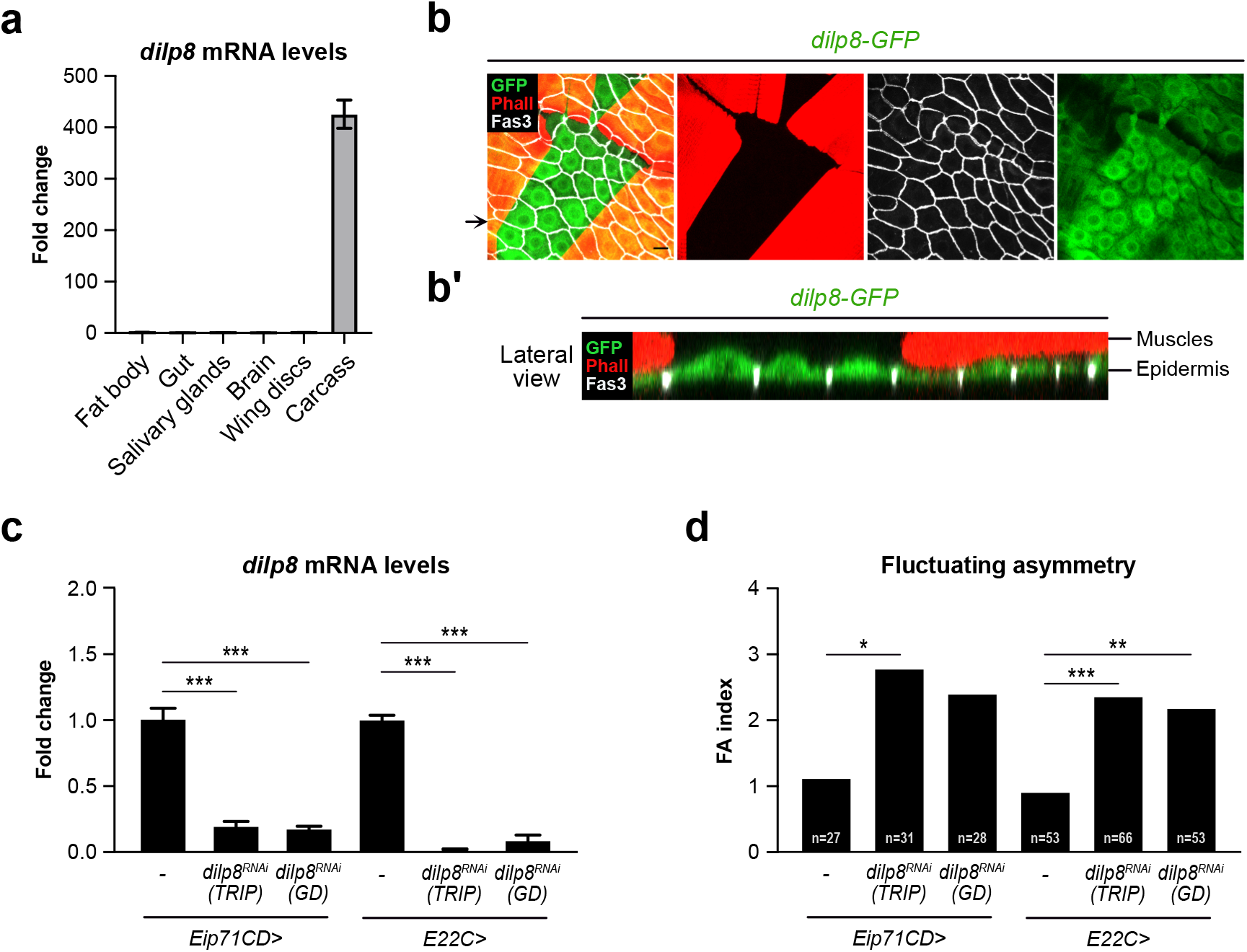
The epidermis is the source of Dilp8 at the WPP stage. (a) Measurement of *dilp8* mRNA levels by qRT-PCR on dissected tissues (*w^1118^* control strain) at the WPP stage. Values are expressed as fold changes relative to the fat body and error bars represent SEM. (b) Maximal projection of a fillet preparation showing expression of the *dilp8-GFP* reporter at the WPP stage. Phalloidin (Phall, in red) staining marks actin filaments in the muscles and Fasciclin 3 (Fas3, in grey) marks the cell membranes of epidermal cells. The arrow on the side indicates the plane for lateral view reconstruction. (b’) Lateral view of the preparation presented in (b), showing that *dilp8-GFP* is expressed in the layer corresponding to epidermal cells and not in the muscles. (c) Measurement of *dilp8* mRNA levels by qRT-PCR on whole animals at the WPP stage upon RNAi-mediated downregulation of *dilp8* in the epidermis. Values are expressed as fold changes relative to controls without RNAi. Error bars represent SEM. *** p<0.001, one-way ANOVA. (d) FA indexes of adult wings upon RNAi-mediated downregulation of *dilp8* in the epidermis. n values indicate the number of pairs analyzed; *** p<0.001, ** p<0.01 and * p<0.05, for *Eip7 1CD>dilp8^RNAi^ GD* p=0.052; F-tests. Experiments in (c) and (d) were done at 29°C. WPP: white prepupa.

Finally, we observed that silencing *dilp8* expression in epidermal cells is sufficient to induce adult wing FA (Fig. 3d), while downregulation of *dilp8* in the muscles does not affect developmental stability (Fig. S2e).

Taken together, our results indicate that the epidermis is the source of a burst of *dilp8* expression at the WPP stage that triggers organ size adjustment.

### A hormonal cross-talk between Ecdysone and Dilp8 defines a time window for size adjustment at the larva-to-pupa transition

The sharp expression of *dilp8* in the WPP epidermis is indicative of a tight spatial and temporal transcriptional control. Temporally, ecdysone titers increase gradually during the L3 stage and reach maximum levels at the WPP stage^18^. Therefore, the peak of *dilp8* expression at WPP could rely on ecdysone. To test this possibility, we silenced expression of the *ecdysone receptor (EcR)* gene specifically in the epidermis using a weak RNAi line to prevent larval or early pupal lethality (*Eip71CD>EcR^RNAi^pan* and *E22C>EcR^RNAi^pan*), and observed a strong decrease in *dilp8* expression at WPP (Fig. 4a). Interestingly, other pathways known to control *dilp8* expression in the context of tissue repair^11,16,17,19^, like Hippo, JNK and Xrp1 signaling, are not required for epidermal *dilp8* expression at the WPP stage (Fig. S3a,b).

**Figure 4.**
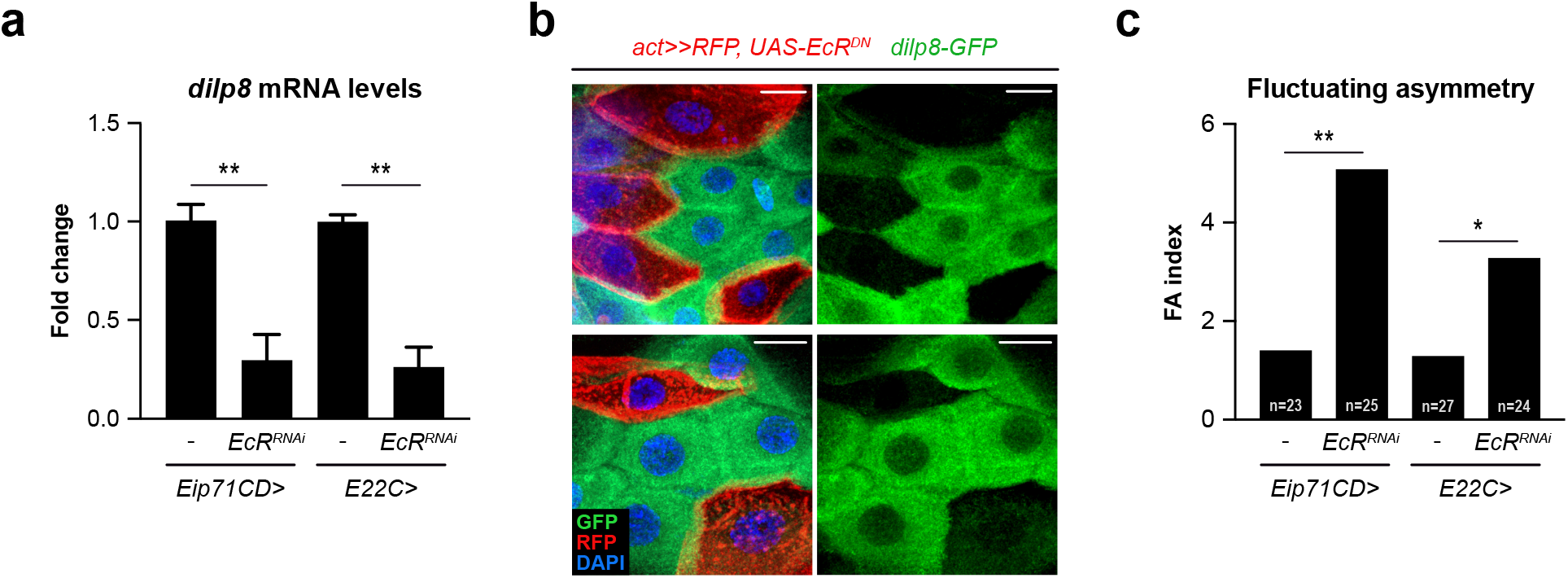
Ecdysone signaling induces *dilp8* expression in the epidermis. (a) Measurement of *dilp8* mRNA levels by qRT-PCR on whole animals at the WPP stage upon RNAi-mediated downregulation of the ecdysone receptor *EcR* in the epidermis. Values are expressed as fold changes relative to controls without RNAi. Error bars represent SEM. ** p<0.01, one-way ANOVA. (b) Two representative examples of epidermis of animals at the WPP stage showing expression of the *dilp8-GFP* reporter in control cells and clones of cells expressing a dominant-negative isoform of EcR (EcR^DN^; cells marked with RFP and shown in red). Scale bars represent 2μm and DAPI stains the nuclei. (c) FA indexes of adult wings upon RNAi-mediated downregulation of *EcR* in the epidermis. n values indicate the number of pairs analyzed; ** p<0.01 and * p<0.05, F-tests. Experiments in (a) and (c) were done at 29°C. WPP: white prepupa.

To confirm these expression results, we analyzed epidermal cells at the WPP stage after clonal expression of a dominant-negative form of EcR (*EcR^DN^*), which binds to the promoter region of target genes but is deficient for transcriptional activation. In *EcR^DN^*-expressing clones, the GFP signal corresponding to the MiMIC *dilp8-GFP* reporter disappears, in contrast with neighboring control cells (Fig. 4b). This confirms the cell-autonomous control of *dilp8* expression by EcR signaling in epidermal cells.

In addition to these expression data, we investigated the role of EcR signaling upstream of Dilp8 in the control of developmental stability. Inhibiting EcR function in the epidermis (*Eip71CD> EcR-RNAi, E22C> EcR-RNAi*) leads to a significant increase in adult wing FA (Fig. 4c). This establishes that a functional cross talk between ecdysone and Dilp8 takes place at WPP in the epidermis for the control of size adjustment.

### Dilp8 controls developmental precision through a feedback on ecdysone signaling in target tissues

In conditions of tissue injury, Dilp8 delays development by inhibiting the peak of ecdysone that triggers the L/P transition^11,12^. To investigate whether Dilp8 also acts upstream of ecdysone for organ size adjustment, we compared the levels of circulating ecdysone in controls and *dilp8^KO/KO^* mutants at several timepoints around the L/P transition. We observed a modification of the peak of ecdysone in *dilp8^KO/KO^* animals, with a significant increase in circulating ecdysone at the WPP stage (0h APF), followed by a sharper decrease between 2h and 8h APF (Fig. 5a). Strikingly, the increase in ecdysone titer occurs precisely when *dilp8* expression normally peaks, suggesting that Dilp8 operates a fast and precise control of the intensity and timing of ecdysone accumulation.

**Figure 5.**
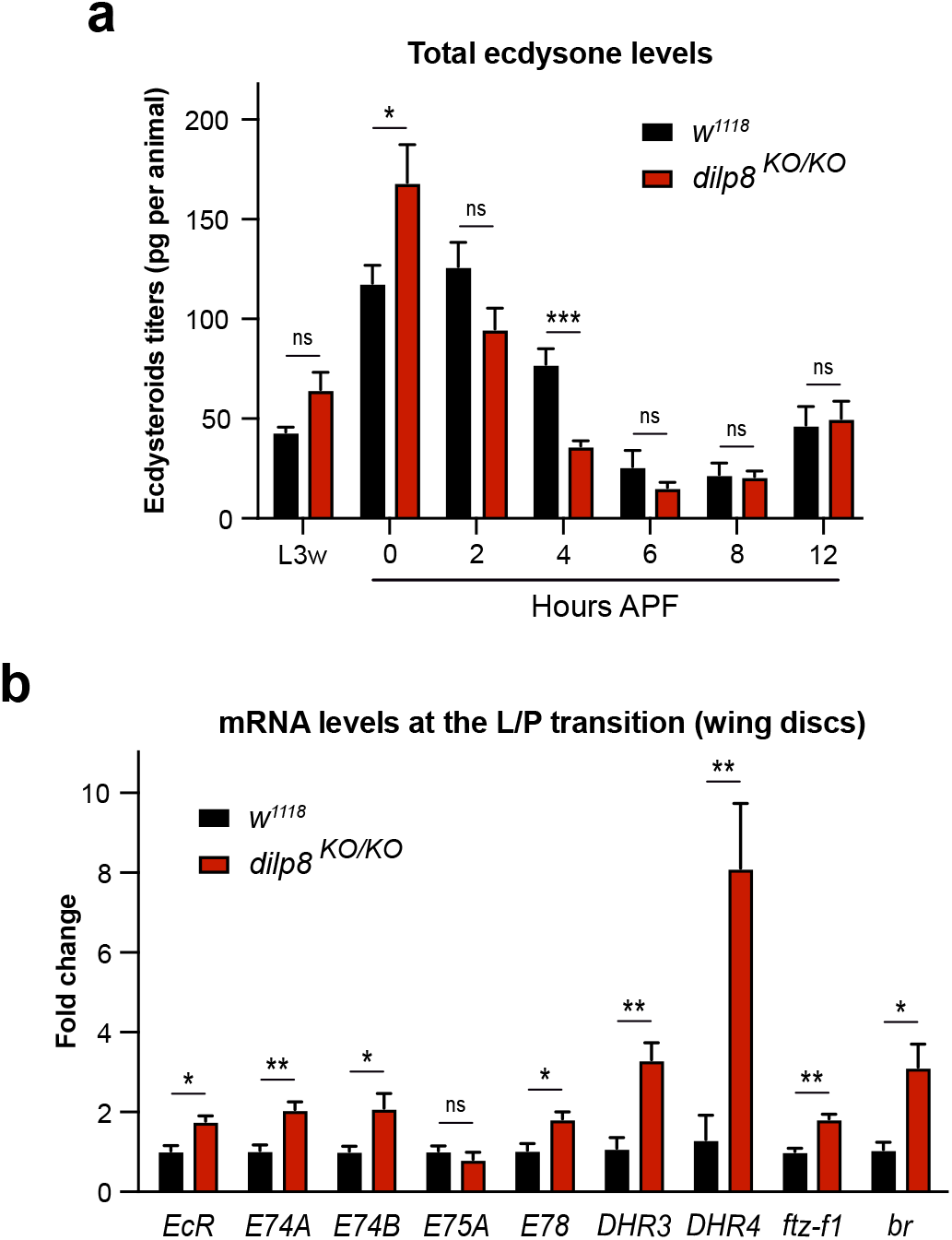
Dilp8 is required to adjust ecdysone levels at the L/P transition. (a) Measurements of ecdysteroids titers in whole animals at the indicated timepoints for controls (in black) and *dilp8^KO/KO^* (in red) mutant animals. At L3w, n=3 independent quantifications were performed per genotype; at 0h AFP, n=7 per genotype; for other time points, n=6 per genotype. Error bars represent SEM. *** p<0.001, * p<0.05 and ns=not significant, t-tests. (b) Measurement of ecdysone targets by qRT-PCR on dissected wing imaginal discs at the WPP stage in controls (in black) and *dilp8^KO/KO^* (in red) mutant animals. Values are expressed as fold changes relative to controls. Error bars represent SEM. ** p<0.01, * p<0.05 and ns=not significant, t-tests. L3w: wandering late L3 stage, APF: after pupa formation, L/P transition: larva-to-pupa transition, WPP: white prepupa.

We next assessed whether the levels of ecdysone signaling in target tissues is modified in the absence of Dilp8 function. For this, we compared the expression levels of *EcR* target genes in dissected wing imaginal discs from *dilp8^KO/KO^* and control animals. We observed that 8 out of 9 *EcR* target genes were significantly upregulated in wing discs in the absence of Dilp8 (Fig. 5b), indicating a clear effect on the intensity of EcR signaling in target tissues at WPP.

Therefore, Dilp8 acts to ensure proper level and timing of circulating ecdysone and, consequently, of EcR signaling in target tissues, allowing developmental adjustment during the critical post-WPP phase.

## DISCUSSION

Precision and stability are fundamental properties of many developmental processes, albeit poorly understood. Paired symmetrical organs have proven useful to quantify stochastic variability of developmental processes. However, studies on how bilateral organ symmetry is established have been limited by the difficulty to precisely quantify 3D morphogenesis on both sides of developing organisms. We provide here the first evaluation of bilateral wing disc development in *Drosophila.*

Unexpectedly, we find that discs adjust their volume through a major adjustment step taking place immediately after the L/P transition. This contrasts with recent descriptions made in zebra fish where symmetrically developing inner ears and somites adjust progressively over time^20,21^. We show that the relaxin-like Dilp8 is required for a major adjustment step taking place after WPP. We also identified a minor adjustment of L-R variability earlier in larval development. However, this adjustment is also observed in *dilp8^KO/KO^* animals, indicating that it relies on a separate mechanism.

These findings contrast with the mechanism by which Dilp8 induces a developmental delay following alterations of disc growth. In that case, Dilp8 is produced by ill-growing tissues and triggers a feedback mechanism on ecdysone production, allowing coupling of the growing state of organs with the major developmental transition at the end of the juvenile period. We show here that, in the absence of perturbation, Dilp8 is produced in a specific tissue, the epidermis, and is required at a precise stage to ensure organ size adjustment. We conclude that Dilp8 participates in a timer mechanism defining a time window for developmental robustness.

We demonstrate a reciprocal feedback between ecdysone and Dilp8 taking place at the WPP stage: while ecdysone is needed for *dilp8* expression, Dilp8 feeds back on ecdysone production and adjusts its levels of signaling in target tissues. We show that none of the upstream signals needed for Dilp8 induction in response to tissue stress (i.e. JNK, Xrp1) is needed for its developmental expression. Although previously shown to contribute to *dilp8* expression^17^, the transcriptional activator Yorkie/Scalloped does not participate in physiological *dilp8* induction at the WPP. Noticeably, removing a Yki/Sd response element in the *dilp8* promoter leads to a rather mild increase in developmental stability^17^. This indicates that Yki/Sd plays a limited role in *dilp8*-dependent developmental stability, distinct from the major regulation step occurring at WPP.

Expression of *dilp8* in the larval epidermis indicates that epidermal cells play a major endocrine role at the larva-to-pupa transition. Intriguingly, while most larval tissues express EcR and Usp and respond to ecdysone during this critical transition, only epidermal cells contribute to EcR-dependent *dilp8* expression. This could result from the specific functional preponderance of the EcR-B2 isoform in the epidermis, compared to other tissues with balanced contributions of several EcR isoforms^22^. Alternatively, specific co-factors present in the epidermis are possibly required together with EcR for *dilp8* induction at WPP. Taiman (Tai) is a co-factor of EcR required for the induction of *dilp8* in wing discs overexpressing Yki^23^, but no significant decrease in *dilp8* expression was observed after silencing *Tai* in epidermal cells (our unpublished data). Therefore, specific epidermal co-factors of EcR required for *dilp8* induction at WPP remain to be identified. The larval epidermis undergoes important transformations at the WPP. A series of contractions shorten the size of the future body and the cuticle sclerotizes to produce a rigid pupal case. These events follow a precisely staged sequence allowing progression into pupal development. Interestingly, epidermally-produced Dilp8 is required for proper accomplishment of this complex behavioral series^24^. Therefore, the production of Dilp8 from the epidermis could allow an integration of major morphological and timer functions needed at the L/P transition.

A parallel should be made between emerging endocrine properties of the larval epidermis presented here and the established neuroendocrine functions of the vertebrate skin. Cutaneous structures respond to, but also generate, a large number of neuromodulators and hormones, which participate in skin homeostatic functions including metabolic activity, tissue repair, immune response (for review^25^). Several neuropeptides were identified from amphibian skin before being found in neural tissues, and human skin recapitulates the TRH/TSH/thyroid and the CRH/ACTH/Cortisol axes found in the central brain^26,27^. These observations have suggested an ancestral function for the epidermis as a neuroendocrine organ. In this context our finding of a Dilp8 epidermal function suggests possible conserved cross-talks between neurohormonal brain and epidermal axes.

We and others previously showed that Dilp8 acts through a limited number of Lgr3-positive neurons in the central lobe region of the larval brain to delay ecdysone production and the L/P transition in response to growth impairment^13^–^15^. Interestingly, depleting Lgr3 in this subpopulation of neurons promotes high adult wing FA, suggesting that Dilp8 controls developmental stability through the same neuronal relay^13^.

Our present findings indicate that the temporal accumulation of ecdysone is modified in *dilp8* loss-of-function conditions. Consistently, higher expression levels of EcR targets are found in wing discs at WPP. Collectively, these and our previous results indicate that epidermal Dilp8 acts on ecdysone accumulation though an Lgr3 central relay and modulates the level and timing of EcR signaling in peripheral tissues for disc size adjustment. Interestingly, Dilp8 is a temporal neuromodulator of ecdysone function, both in the context of growth impairment and during normal development. The sharp induction of *dilp8* at WPP and the immediate response on ecdysone levels observed upon *dilp8* loss-of-function (see Fig. 5a,b) indicate that Dilp8-mediated neuro-hormonal action has high temporal definition, an important property for its function as a developmental timer.

In conclusion, our results define a hormonal cross-talk between ecdysone and Dilp8 that is key for developmental precision. This cross-talk has two functions: (i) it defines a time window after larval development during which wing discs adjust their size; (ii) it fine-tunes the levels of ecdysone signaling in the discs, which appears crucial for their size adjustment. The dynamics of ecdysone levels in early pupal wing discs controls a cascade of transcription leading to two waves of cell division/cell cycle exit^28^. Further work will be needed to understand how the systemic ecdysone signal contributes to adjusting organ size during pupal development.

## Methods

### Fly strains and food

The following RNAi lines were obtained from the Vienna Drosophila RNAi Center (VDRC): *UAS-yki^RNAi^* (KK 104523), *UAS-sd^RNAi^* (KK 108877) and *UAS-dilp8^RNAi^* GD (GD 9420). The *w^1118^* (BL 3605), *UAS-GFP* (BL 35786), *UAS-dilp8^RNAi^* TRIP (BL 80436), *dilp8-GFP* (dilp8^MI00727^; BL 33079), *tub-GAL80^ts^:tub-GAL4* (BL 86328), *mef2-Gal4* (BL 27390), *mhc-GAL4* (BL 84298), *UAS-xrp1^RNAi^* (BL 34521), *UAS-bsk^DN^* (BL 6409), *UAS-EcR^DN^* (BL 6869), *UAS-EcR^RNAi^ pan* (BL 29374), *UAS-jub^RNAi^* (BL 32923), *Eip71CD-GAL4* (BL 6871) lines were provided by the Bloomington Drosophila Stock Center. Other lines used in this study were: *nub-GAL4^29^; dilp8^KO^* and *dilp8-full-prom-lacZ^17^*, *E22C-GAL4^30^; hs-Flp, act-FRT-STOP-FRT-GAL4, tub-GAL80^ts^ UAS-RFP* (gift from the Bellaïche Lab).

Animals were reared at 25□°C (unless otherwise stated in the figure legends) on fly food containing, per liter: 14□g inactivated yeast powder, 69 g corn meal, 7.5 g agar, 52 g white sugar and 1.4□g Methyl 4-hydroxybenzoate.

### Temperature-shift (TS) experiments

Crosses of the *UAS-dilp8^RNAi^* line or the control *UAS-GFP* line with the *tub-GAL4, tub-GAL80^ts^* line were left for oviposition during 4h on plates made of 2% agar and 2% sucrose in PBS. The next day, synchronized L1-stage animals were transferred from the agar plates to vials with fly food and kept at 18°C to repress GAL4 activity until the indicated times. At this point, the tubes with synchronized larvae were shifted to 29°C to allow GAL4 activity. For TS at the early L3 stage, animals were transferred to 29° for 24h at 6 days AED. For TS at the WPP stage, animals were transferred to 29° for 24h at 9 days AED, after which most animals had pupariated (remaining larvae were removed).

### Clonal analysis

Crosses of the *hs-Flp, act-FRT-STOP-FRT-Gal4, tub-Gal80^ts^; UAS-RFP* line with a *UAS-EcR^DN^;dilp8-GFP* line were performed in tubes with fly food and left at 25 °C. At L1 stage, a 30 min heat-shock was performed in a 42 °C water bath, using an immersion circulator (Julabo), to provoke a random flip-out activation of GAL4 expression. After the heat-shock, tubes were immediately transferred to 18 °C to repress GAL4 activity and avoid deleterious effects of EcR^DN^ over-expression. At the L3w stage, larvae were shifted to 29 °C for one day to allow maximum activation of EcR^DN^ expression and samples were dissected at the WPP stage.

### Quantitative RT-PCR

Larvae or pupae were collected at the indicated stages. Whole animals or dissected tissues were frozen in liquid nitrogen. Total RNA was extracted using a QIAGEN RNeasy Lipid Tissue Mini Kit (for whole larvae samples, processed with QIAcube after homogenization with the Tissue Lyser II (Qiagen)) or a QIAGEN RNeasy Micro Kit (for dissected wing discs) according to the manufacturer’s protocol. RNA samples (2-3μg per reaction) were treated with DNase when necessary and reverse-transcribed using SuperScript II reverse transcriptase (Invitrogen), and the generated cDNAs were used for real-time PCR (StepOne Plus, Applied Biosystems) using Power SYBR Green PCR mastermix (Applied Biosystems). Samples were normalized to *rp49* and fold changes were calculated using the ΔΔCt method; P values are the result of t-tests or ANOVA tests provided by Graphpad. At least three separate biological samples (5-10 animals each) were collected for each experiment and triplicate measurements were performed. The following primers were used:

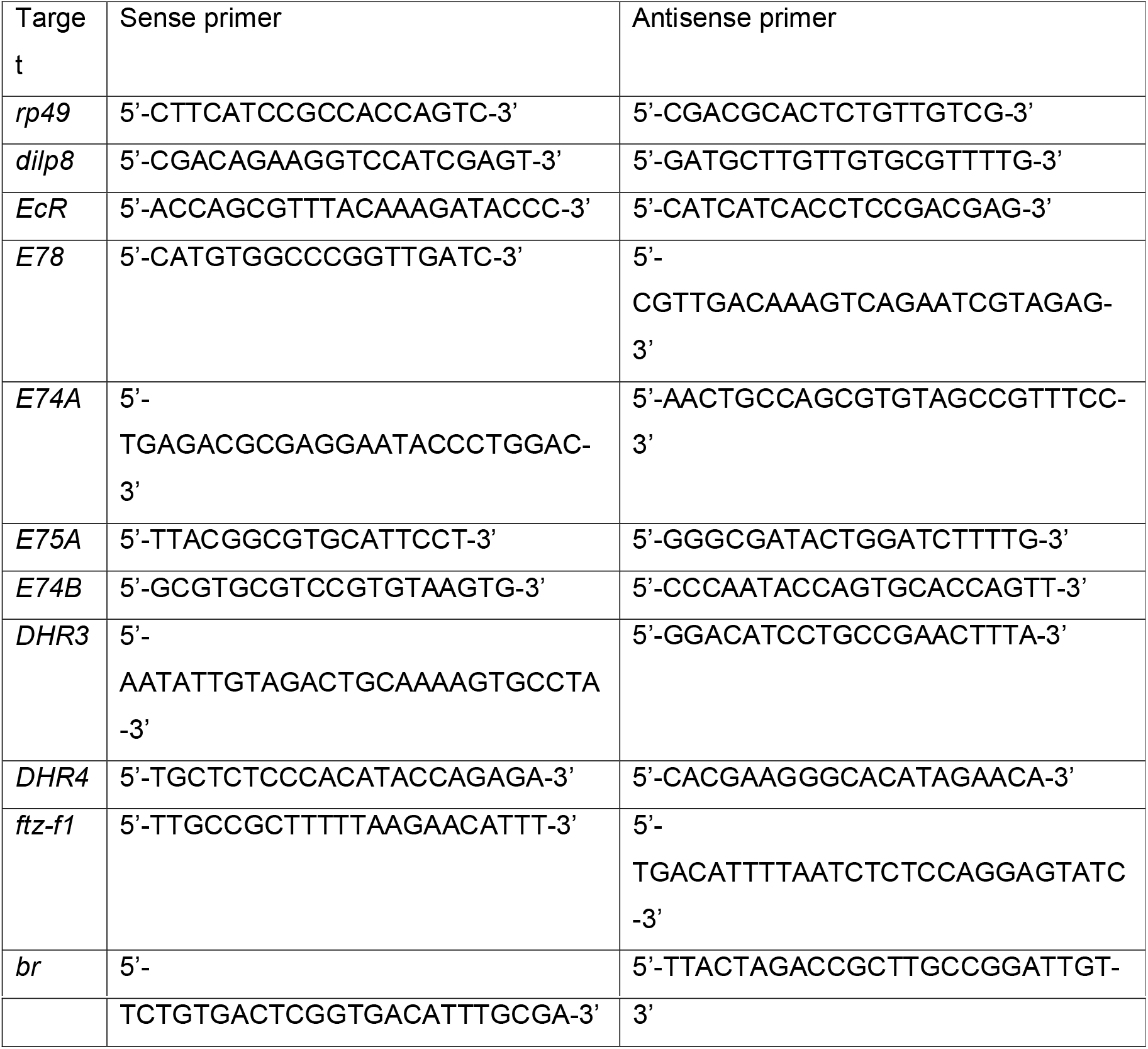

### Measurement of the FA index

In the case of wing primordia measurements (Fig. 1), animals were dissected at the given timepoints, fixed in 4% formaldehyde (Sigma) in PBS for 30□min at room temperature, and washed in PBS. For early pupae (7h APF), animals were dissected as described in ^31^. The left- and right-wing discs of each individual were mounted without coverslip on “Cellview” cell culture dishes with glass bottom (Greiner Bio-one), in order to preserve the original structure volume. Imaging was performed with a Zeiss LSM900 Inverted Laser Scanning Confocal Microscope using exactly the same settings for each pair. The confocal Z-stacks were processed with the Imaris software using exactly the same settings for each pair, and surfaces of the *nub>GFP* signal were generated to faithfully represent the original structure volume.

In the case of adult wing measurements, adult flies of the appropriate genotypes were collected, stored in ethanol and mounted in a lactic acid:ethanol (6:5) solution. Wings were dissected and mounted in pairs. Pictures were acquired using a Leica Fluorescence Stereomicroscope MZ16 FA with a Leica digital camera DFC 490.

We used the FA index (FAi) number 6 as described by Palmer and Strobeck^2^ to assess intraindividual size variations between left and right wing primordia or adult wings:

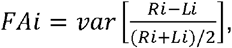

where Ri and Li are the sizes of the right (R) and left (L) dissected discs or adult wings of the same individual. This FA index was chosen because it normalizes left-right differences to average tissue size, and therefore prevents biases linked to experimental effects on average size (such as temperature changes or developmental timepoints). Figures represent the FAi x10^4^; P values are the results of a F test. Only females (both for dissected discs and for adult wings) were analyzed.

### Immunostainings of larval tissues

Tissues dissected from larvae or pupae in 1X PBS at the indicated stages were fixed in 4% formaldehyde (Sigma) in PBS for 30 min at room temperature, washed in PBS containing 0.3 % Triton-X-100 (PBT), blocked in PBT containing 2% BSA and incubated overnight with primary antibodies at 4°C. The next day, tissues were washed, blocked again and incubated with secondary antibodies at 1/250 dilution (Alexa Fluor 647, 555 and 488 from Jackson ImmunoResearch) and/or 1/100 dilution of Alexa Fluor 647 Phalloidin (ThermoFisher) for 2□h at room temperature. Samples were mounted in Vectashield (Vector Labs) or SlowFade Diamond with DAPI (ThermoFisher). Fluorescence images were acquired using a Zeiss LSM900 Inverted Laser Scanning Confocal Microscope and processed using Image J.

The following primary antibodies were used: chicken anti-GFP, 1/10000 (Abcam), mouse anti-FasIII, 1/50 (Developmental Studies Hybridoma Bank) and mouse anti-beta-galactosidase, 1/200 (Promega). The epidermis of WPP was dissected following fillet preparation protocols described in ^32^.

### Ecdysteroids extraction and quantification

For ecdysteroids extraction, 6 to 10 whole animals at the indicated stages were collected for each biological replicate in 2ml Eppendorf tubes, frozen in liquid nitrogen and stored at −80°C. Samples were homogenized using a metal bead and Tissue Lyser II (Qiagen) in 0.3ml of methanol, centrifuged at maximum speed for 5 min at room temperature (RT) and the supernatant was transferred to a new tube. 0.3ml of methanol was added, and samples were mixed using Vortex. This procedure was repeated using 0.3ml of ethanol, so that the pooled samples contained a total volume of 0.9ml, and were stored at −80°C.

For quantification, the extracted samples were centrifuged at maximum speed for 5 min (RT) to remove any remaining debris and divided into two tubes to generate technical replicates. The cleared samples were evaporated using a Speedvac centrifuge equipped with a cold trap. The following steps were performed using the 20-Hydroxyecdysone ELISA kit (Bertin Bioreagent), with the following modifications: After evaporation, the precipitate was redissolved in 200μl of ELISA buffer (EIA Buffer). It was critical to aid the re-dissolution of the precipitate by scraping it with a pestle and by vigorous vortexing, until no more was visible on the walls of the tubes. The ELISA plates were loaded with samples and a standard curve as indicated by the manufacturer, incubated overnight at 4°C and read with a Microplate Reader (Tecan Sunrise) at 405 nm. Data analysis was performed as indicated in the Bertin 20-Hydroxyecdysone ELISA kit manual.

## Statistics

For comparison of the means, two-tailed T-tests or ANOVA analysis (as indicated in the figure legends) were performed using GraphPad. For comparisons of FAi values, F-tests provided by Microsoft Excel were used. In all cases, n values are indicated for each experiment in the corresponding figures or figure legends.

## DATA AVAILABILITY

The data that support the findings of this study are available from the corresponding authors upon reasonable request.

## ACKNOWLEDGMENTS

We thank David Lubensky, Ojan Khatib-Damavandi and Marie-Anne Félix for insightful discussions; Paula Santabárbara-Ruiz for help with the experiments and members of the laboratory for discussions and comments on the manuscript; Julio Lopes Sampaio for help with ecdysteroids extractions; the Bloomington Stock Center and the Vienna Drosophila RNAi Center for fly stocks; the PICT-IBiSA@BDD light-microscopy facility of Institut Curie. This work was supported by Institut Curie, CNRS, INSERM, FRM, European Research Council (Advanced Grant n°694677 to P.L.), Labex DEEP program (ANR-11-LABX-0044, ANR-10-IDEX-0001-02), PSL (PhD fellowship to K.E.) and the Marie Sklodowska-Curie Actions (fellowship n° 897309 to D.B-O).

## AUTHORS CONTRIBUTIONS

Conceptualization and methodology: L.B., D.B-O., D.S.A., J.C., P.L. Investigations: L.B., D.B-O., K.E.M, F.B., D.S.A., J.C., S.N. Writing: L.B., D.B-O., P.L. Funding acquisition and supervision: P.L.

## COMPETING INTERESTS

The authors declare no competing financial interests.

## MATERIALS AND CORRESPONDENCE

Correspondence and material requests should be addressed to.

## Notes

### Competing Interest Statement

The authors have declared no competing interest.

### Summary of Updates

The control of organ size mainly relies on precise autonomous growth programs. However, organ development is subject to random variations, called developmental noise, best revealed by the fluctuating asymmetry observed between bilateral organs. The developmental mechanisms ensuring bilateral symmetry in organ size are mostly unknown. In Drosophila, null mutations for the relaxin-like hormone Dilp8 increase wing fluctuating asymmetry, suggesting that Dilp8 plays a role in buffering developmental noise. Here we show that size adjustment of the wing primordia involves a peak of Dilp8 expression that takes place sharply at the end of juvenile growth. Wing size adjustment relies on a cross-organ communication involving the epidermis as the source of Dilp8. We identify ecdysone signaling as both the trigger for epidermal dilp8 expression and its downstream target in the wing primordia, thereby establishing reciprocal feedback between the two hormones as a systemic mechanism controlling organ size precision. Our results reveal a hormone-based time window ensuring fine-tuning of organ size and bilateral symmetry.

