## Supplementary figures for "Dilp8 controls a time window for tissue size adjustment in *Drosophila*"

### Figure S1

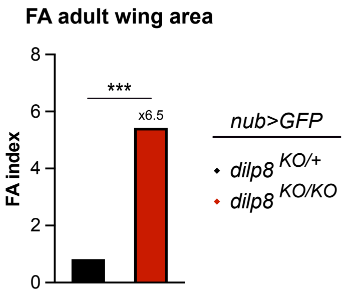


FA indexes in adults of the same genotypes and issued from the same crosses as presented in Figure 1. n=24 pairs of wings for *dilp8*$^{KO/+}$*, nub>GFP* controls and n=19 for *dilp8*$^{KO/KO}$*, nub>GFP* animals. *** $p<0.001$, F-test.

### Figure S2

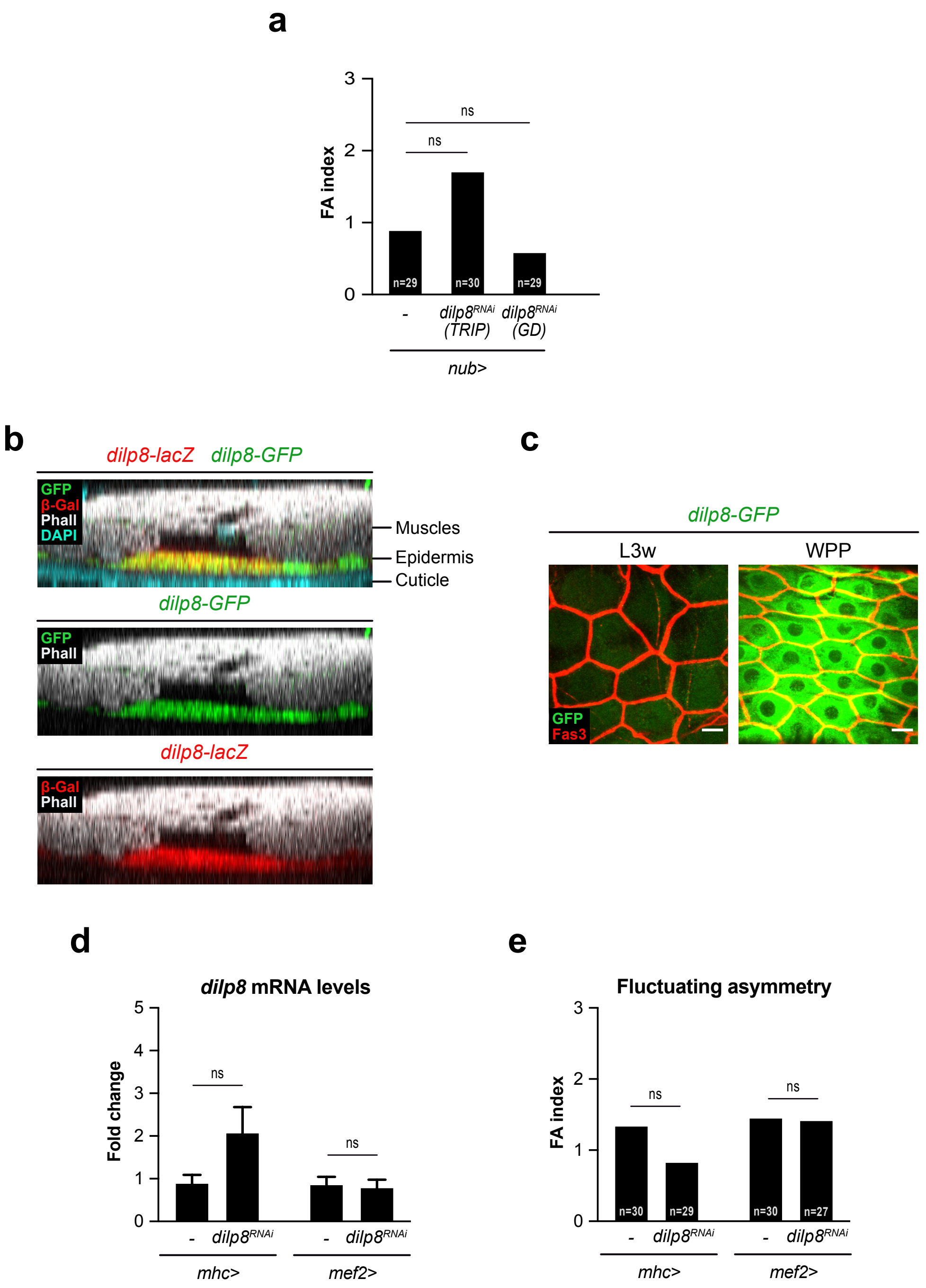


**(a)** FA indexes of adult wings upon RNAi-mediated downregulation of *dilp8* in the wing imaginal discs using *nub-GAL4*. n values indicate the number of pairs analyzed; ns=not significant, F-test. **(b)** Lateral view of a fillet preparation at the WPP stage showing expression of the *dilp8-lacZ* (in red) and *dilp8-GFP* (in green) transcriptional reporters in the epidermis. Phalloidin (Phall) staining marks the muscles and DAPI is used here to stain the cuticle. **(c)** Maximal projections of fillet preparations showing expression of the *dilp8-GFP* reporter at the wandering L3 stage (L3w) and at the WPP stage. Fasciclin 3 (Fas 3, in red) marks the outer membranes of epidermal cells. Scale bars represent 20 microns. **(d)** Measurement of *dilp8* mRNA levels by qRT-PCR on whole animals at the WPP stage upon RNAi-mediated downregulation of *dilp8* (*UAS-dilp8*$^{RNAi}$ TRIP line) in the muscles. Values are expressed as fold changes relative to controls without RNAi. Error bars represent SEM. ns=not significant, one-way ANOVA. **(e)** FA indexes of adult wings upon RNAi-mediated downregulation of *dilp8* (*UAS-dilp8*$^{RNAi}$ TRIP line) in the muscles. n values indicate the number of pairs analyzed; ns=not significant, F-test. Experiments in (a, d, e) were done at 29°C.

### Figure S3

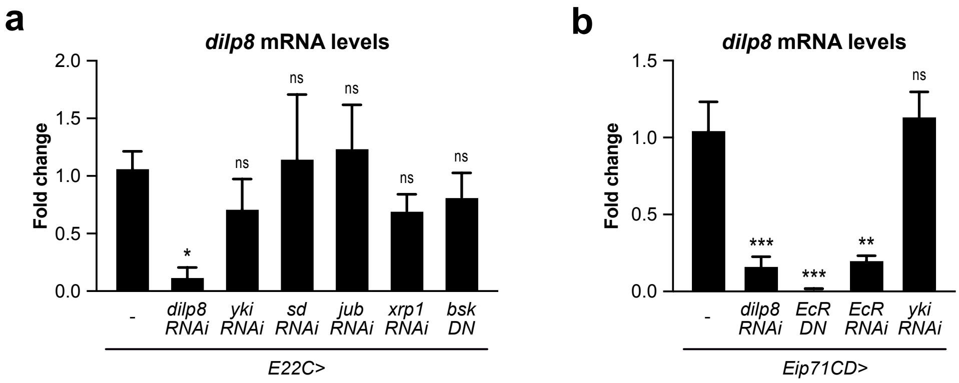


**(a)** and **(b)** Measurement of *dilp8* mRNA levels by qRT-PCR on whole animals of the indicated genotypes at the WPP stage. Values are expressed as fold changes relative to controls without RNAi. Error bars represent SEM. *** p<0.001, * p<0.05 and ns=not significant, one-way ANOVA. Experiments were done at 25°C.
